# A short sequence targets transmembrane proteins to primary cilia

**DOI:** 10.1101/2024.04.02.587776

**Authors:** Viviana Macarelli, Edward Harding, David C. Gershlick, Florian T. Merkle

## Abstract

Primary cilia are finger-like sensory organelles that extend from the bodies of most cell types and have a distinct lipid and protein composition from the plasma membrane. This partitioning is maintained by a diffusion barrier that restricts the entry of non-ciliary proteins, and allows the selective entry of proteins harboring a ciliary targeting sequence (CTS). However, CTSs are not stereotyped, and previously-reported sequences are insufficient to drive efficient ciliary localization across diverse cell types. Here, we describe a short peptide sequence that efficiently targets transmembrane proteins to primary cilia in all tested cell types, including human neurons. We generate human induced pluripotent stem cell (hiPSC) lines stably expressing a transmembrane construct bearing an extracellular HaloTag and intracellular fluorescent protein, that enables bright, specific labeling of primary cilia in neurons and other cell types. We demonstrate the utility of this resource by developing an image analysis pipeline for the automated measurement of primary cilia to detect changes in their length.

## Introduction

Primary cilia are non-motile structures that protrude from the plasma membrane of most eukaryotic cell types. While their exact structure and protein composition varies by cell type, it is widely accepted that they play key sensory roles (Anvarian et al. 2019). For example, photoreceptor outer segments are a type of specialized primary cilium, olfactory sensory neurons utilize primary cilia to efficiently detect odorants, and primary cilia in hypothalamic neurons likely sense neuropeptides critical for normal body weight regulation (Y. Wang et al. 2021; Davenport et al. 2007; Brewer et al. 2022). This sensory specialization is mediated by the ciliary localization and concentration of certain ion channels, receptor tyrosine kinases, G-protein coupled receptors (GPCRs), and their downstream mediators (Anvarian et al. 2019). The close physical proximity of these proteins in the primary cilium enables efficient signal transduction, and its small volume can lead to dramatic changes in the concentration of second messengers such as cAMP or Ca^2+^ (Nachury and Mick 2019).

Selective ciliary protein localization is regulated by the distinct structural features of primary cilia, that are organized around a central microtubule-based structure known as the axoneme. Ciliary proteins and cargo vesicles are actively transported up and down the axoneme by kinesin and dynein molecular motors, respectively, using intraflagellar transport complexes A (IFT-A) and B (ITF-B). The IFT machinery assembles into larger macromolecular structures known as ‘IFT trains’ that can be observed along the axoneme or at the ciliary base awaiting entry into the cilium (Nakayama and Katoh 2020; van den Hoek et al. 2022). Many proteins destined for primary cilia harbor a ciliary targeting or localisation sequence (CTS). In contrast to signal peptides that determine canonical translocation with well-characterized sequences, CTS sequences are diverse and their interaction partners and pathways for ciliary targeting are not well understood (Nachury, Seeley, and Jin 2010; Garcia-Gonzalo and Reiter 2012). Indeed, some ciliary proteins appear to bypass the Golgi altogether (Witzgall 2018).

Once at the cilium, ciliary entry is regulated by the transition zone (TZ) at the proximal end of the axoneme near its connection to the basal body, where it is anchored to the ciliary membrane via Y-link protein complexes. The TZ forms a selectively permeable diffusion barrier for both proteins and lipids, effectively separating the ciliary membrane from the rest of the cell (Gonçalves and Pelletier 2017). The ciliary entry of proteins >∼100 kDa and vesicles containing membrane-associated proteins is regulated by protein complexes named after ciliopathies resulting from their functional disruption (Long and Huang 2019; Nachury and Mick 2019; Hu et al. 2010; Jensen and Leroux 2017; Kee et al. 2012);.

Experimentally targeting proteins to primary cilia is a powerful way to study ciliary structure and function in live cells; to characterize the ciliary proteome or potentially revealing new therapeutic targets that act in primary cilia. Others have shown that fluorescent reporters localize to primary cilia when fused to the coding sequences of somatostatin receptor 3 (SSTR3) (O’Connor et al. 2013; Berbari, Johnson, et al. 2008; Guo et al. 2019), the serotonin receptor 6 (5HT6), or ADP ribosylation factor-like GTPase 13B (ARL13B) (Jiang et al. 2019; Moore et al. 2016). The same is true for the first 203 residues of nephronophthisis 3 (NPHP3), which contains an N-terminal myristoylation site and an N-terminal coiled-coil domain required for proper ciliary targeting (Nakata et al. 2012), and is sufficient to target fusion proteins to primary cilia in some cell types (Mick et al. 2015). However, existing ciliary targeting constructs including ARL13B-GFP and N-terminal NPHP3-GFP do not localize to cilia in some cells derived from human induced pluripotent stem cells (hiPSCs), limiting opportunities to study cilia in diverse human cell types. In addition, even in cases where off-target expression of these constructs is not an issue, overexpressing whole functional proteins or protein domains may inadvertently alter normal ciliary function, making interpretation challenging. An ideal CTS would facilitate specific ciliary localisation in many cell types, be compact in order to preserve protein function, and not perturb the endogenous function of the primary cilium.

We set out to find new methods to specifically label the primary cilia of human iPSC-derived neurons, including cortical neurons and hypothalamic neurons reported to be enriched in obesity-associated GPCRs, such as the melanocortin 4 receptor (MC4R) (Y. Wang et al. 2021; Loktev and Jackson 2013) We therefore generated cytoplasmic or membrane-localized reporter constructs and tested the ability of candidate CTSs to mediate ciliary targeting. We identified two sets of CTSs, one from the melanin concentrating hormone receptor 1 (MCHR1), and one from polycystin 2 (PC2/PKD2) sufficient to target a transmembrane construct to primary cilia in hPSC-derived neurons and all three other commonly-used ciliated cell lines we tested. We therefore generated stable iPSC lines expressing a construct with or without the 3x CTS array, which labels either the primary cilium or plasma membrane, respectively, in hiPSC-derived cell types. We demonstrate the utility of this tool by testing the effect of compounds that alter ciliary length in live human neurons, and provide the reagents we established as a resource to the community.

## Results

### Identification of CTS sufficient for ciliary targeting in human neurons

We set out to study the function of primary cilia in live human cortical and hypothalamic neurons derived from hiPSCs (Fig. 1A, S1A,B). To confirm that these neurons are indeed ciliated, we first differentiated the ‘reference’ hiPSC line KOLF2.1J (Pantazis et al. 2022) into hypothalamic neurons as previously described (Merkle et al. 2015; H.-J. C. Chen et al. 2023), and immunostained these cultures for the neuronal marker microtubule associated protein 2 (MAP2) and the primary cilia markers ADP ribosylation factor like GTPase 13B (ARL13B) and adenylate cyclase 3 (ADCY3). We found that hiPSC-derived hypothalamic neurons, including appetite-suppressing proopiomelanocortin (POMC) neurons, had cilia-shaped structures about 3-6 microns in length that projected from neuronal cell bodies, and were immunopositive for primary cilia markers (Fig. 1B,C). Next, we set out to label these neuronal cilia with existing fusion constructs of fluorescent reporters and protein domains reported to mediate ciliary targeting. We therefore introduced the Cilia-APEX (Mick et al. 2015) expression plasmid, in which ciliary localisation is mediated by the N-terminal NPHP3 CTS, into hiPSC-derived neurons and the ciliated IMCD3 cell line by transient transfection. While we confirmed its ciliary localization in IMCD3 cells, the construct failed to localize to primary cilia in hiPSC-derived human neurons across a broad range of tested plasmid concentrations, transfection methods, or neuronal maturation time points (Fig. S1C,D).

**Figure 1:**
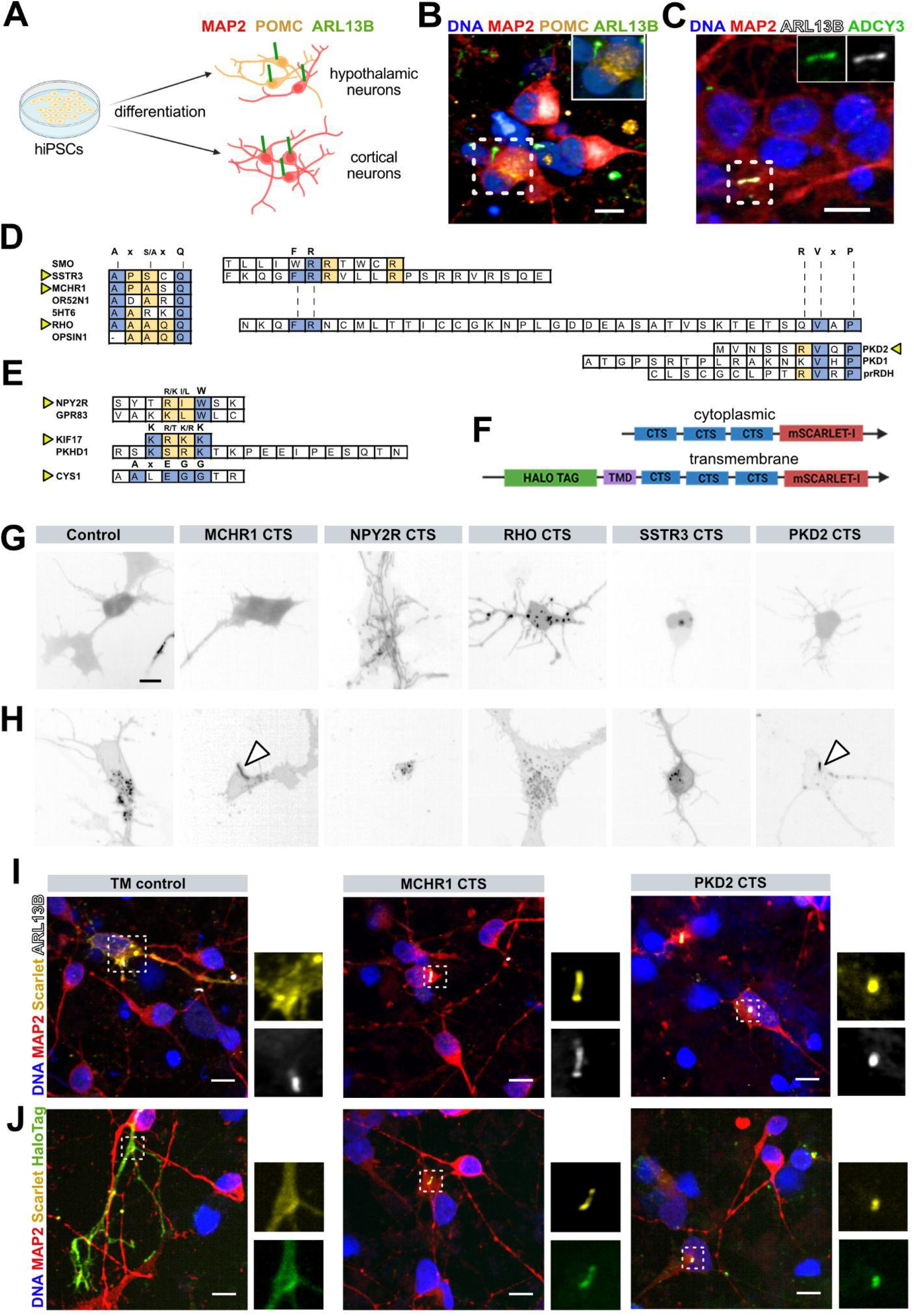
Targeting of transmembrane constructs to primary cilia in human neurons. **A)** Schematic showing the differentiation of hiPSC into hypothalamic neurons (including POMC) and cortical neurons that have primary cilia expressing the cilia-enriched markers ARL13B and ADCY3. **B)** ARL13B+ primary cilia project from the cell bodies of hypothalamic neurons, including those immunopositive for POMC. **C)** Cortical neuron example cilia co-expressing ARL13B and ADCY3 (also seen in hypothalamic neurons). **D)** Sequences shown to be necessary for ciliary targeting from different proteins, where the consensus sequence is in bold, conserved residues are in blue and less conserved residues are in yellow. Sequences selected in this study are indicated with arrowheads. **E)** As in (D) but for distinct CTSs. **F)** Schematic of construct design, where a 3x array of candidate CTSs followed by mScarlet with or without an N-terminal signal peptide, HaloTag, and transmembrane domain were used to test ciliary targeting. **G,H)** Representative images of mScarlet signal from transiently transfected neurons showing the relative lack of ciliary targeting with cytoplasmic constructs (G), but ciliary localisation with CTSs derived from MCHR1 and PKD2 in transmembrane constructs (H, arrowheads). **I)** Sequences from MCHR1 and PKD2 drove localization of mScarlet to structures immunopositive for ARL13B, and could also be visualized with fluorescent HaloTag ligands (J). Insets show cilia at higher magnification. Scale bars represent 10 µm in B,C,J, G and H.

We hypothesized that this lack of ciliary labeling in human neurons was due to different cell type-specific mechanisms regulating protein trafficking to the ciliary base, or mechanisms regulating ciliary entry. Upon reviewing the literature, we found certain motifs that were recurrently described for ciliary targeting in a certain cell types and protein contexts, including the Ax[S/A]xQ (Berbari, Johnson, et al. 2008; Follit et al. 2010), VxP (J. Wang and Deretic 2014; Ward et al. 2011; Geng et al. 2006; Luo et al. 2004) and FR (Ward et al. 2011; Chadha, Paniagua, and Williams 2021) motifs, and the less characterized [R/K][I/L]W (Loktev and Jackson 2013), KRKK (Dishinger et al. 2010), KTRK (Loktev and Jackson 2013; Follit et al. 2010), and AxEGG (Tao et al. 2009) motifs (Fig1. D,E). We selected CTSs from seven genes to test the ability of each of these motif types to mediate ciliary targeting in human neurons (Table 1).

**Table 1.**
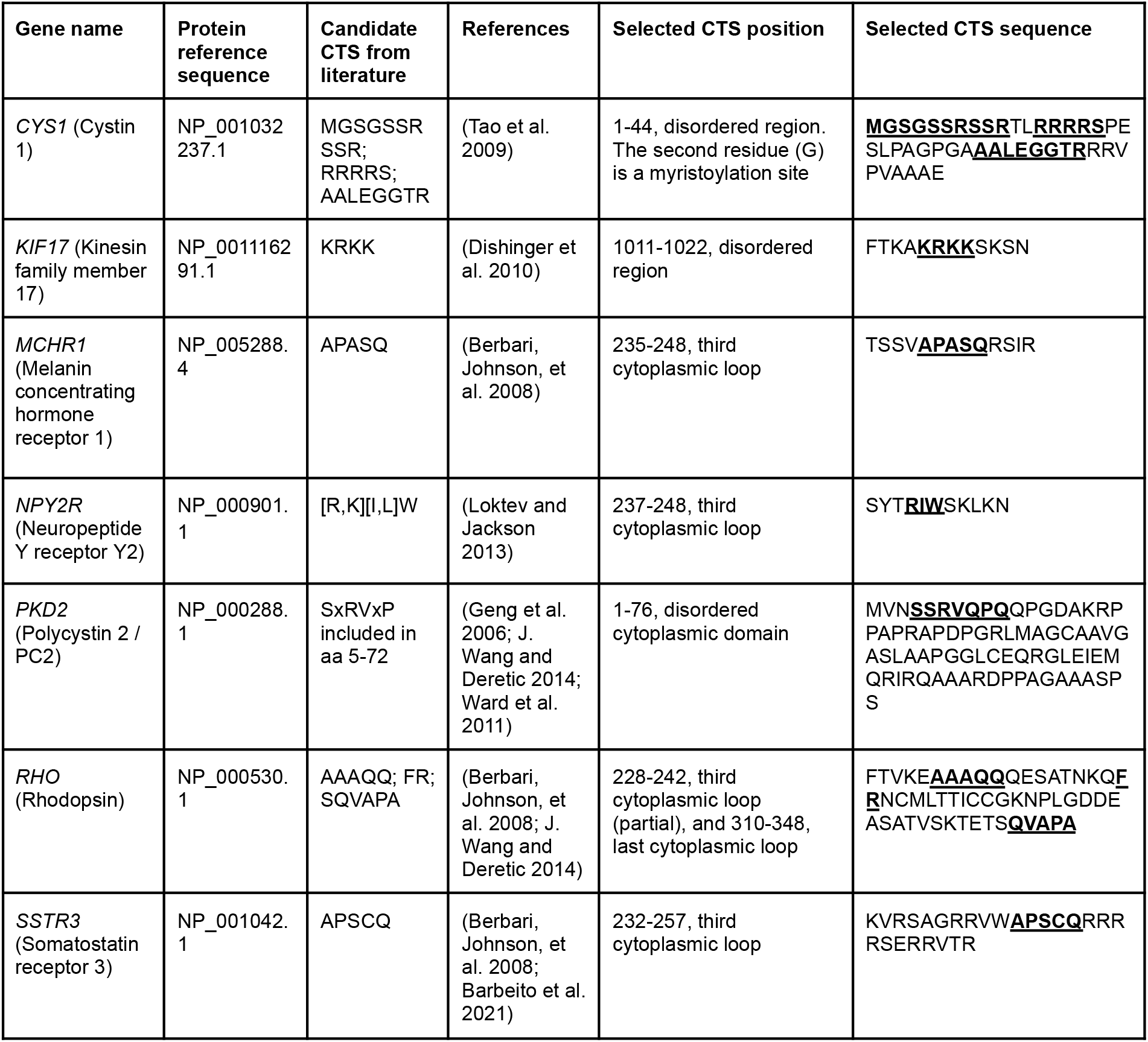
Candidate ciliary targeting sequences (CTS) from candidate genes include core motif sequences (bold and underlined) and flanking residues. Selected sequences (right-hand column) were cloned into a 3x array for testing in hiPSC-derived neurons and other ciliated cell types.

Specifically, we selected regions containing the Ax[S/A]xQ from MCHR1 and SSTR3 (Barbeito et al. 2021) and regions containing the [R/K][I/L]W motif from NPY2R, since all three of these GPCRs localize to primary cilia in hypothalamic neurons (Schou, Pedersen, and Christensen 2015). We also tested candidate CTSs containing a VxP motif from PKD2, and a CTS from RHO that contains three motifs (Ax[S/A]xQ, FR and VxP). Finally, the CTS motifs KRKK and AxEGG were selected from KIF17 and CYS1. We did not consider candidate CTSs from 5HT6 and PKHD1 due to their lower sequence homology between human and mice (Berbari, Johnson, et al. 2008; Follit et al. 2010). We selected 3-5 amino acids on either side of the described core CTS motif, and reasoned that a 3x tandem array of these candidate sequences separated by a short flexible linker (GGGGS) would increase the likelihood of engagement with ciliary localisation machinery (X. Chen, Zaro, and Shen 2013). We omitted the linkers for the CTS from PKD2 due to its relatively large size.

We then cloned these CTS arrays upstream of the bright monomeric red fluorescent protein mScarlet-I to enable visualization of the construct in live cells (Snapp 2005; Bindels et al. 2017). Since ciliary targeting mechanisms are likely distinct for cytosolic and transmembrane or lipid-associated proteins, and it is unclear how isolated CTSs taken from different proteins would act in the context of fusion reporter construct, we generated both cytoplasmic and transmembrane versions of most of these constructs (Garcia-Gonzalo and Reiter 2012) except CYS1 and KIF17. To generate the transmembrane versions, we included an N-terminal IL2 signal peptide and HaloTag to facilitate extracellular fluorescent labeling (Los et al. 2008), a GGGS linker and synthetic transmembrane domain consisting of 22 leucine residues and four basic residues to stabilize the protein in the membrane and promote export through the biosynthetic secretory pathway (Sharpe, Stevens, and Munro 2010; H. Chen and Kendall 1995; Munro 1995), and GS residues just upstream of the CTS array to localize putative ciliary targeting sequences close to the plasma membrane (Fig. 1F, Table 1). We then electroporated these cytoplasmic and transmembrane constructs into hiPSC-derived hypothalamic neurons and screened for the localization of mScarlet cilia-like structures. We found that none of the cytoplasmic constructs we tested showed apparent ciliary enrichment (Fig. 1G, S1D), but that two of the transmembrane constructs containing CTSs derived from MCHR1 or PKD2 localized preferentially to cilia-like structures in transfected cells (Fig. 1H).

To confirm these findings, we fixed and immunostained cultures for neuronal and ciliary markers. We found that while mScarlet was localized to the plasma membrane for transmembrane control constructs lacking CTS arrays (TM control), mScarlet co-localized with ARL13B in cells transfected with constructs containing MCHR1- or PKD2-derived CTSs. Since these constructs also contain an extracellular HaloTag, we added fluorescent HaloTag ligands and found that this fluorescent signal also co-localised with both mScarlet and ciliary markers (Fig. 1I,J). Together, these findings indicate that primary cilia in live human neurons can be visualized with both an intracellular fluorescent protein and a variety of extracellular HaloTag ligands.

### Ciliary targeting across diverse ciliated cell lines

Since the CTS arrays we tested from MCHR1 and PKD2 were sufficient to drive ciliary localisation in iPSC-derived neurons, we next wondered whether constructs containing these sequences would localize to cilia in other cell lines, and how their performance would compare to the ‘Cilia-APEX’ construct encoding a fusion of the first 203 residues of NPHP3 (Nakata et al. 2012), GFP, and the ascorbate peroxidase APEX to facilitate ciliary proximity proteomics (Hung et al. 2016; Mick et al. 2015; May et al. 2021) (Fig. S2A). To replicate published studies, we first confirmed the ciliary targeting of this construct following transient transfection in serum-starved inner medullary collecting duct epithelial (IMCD3), mouse embryonic fibroblasts (NIH-3T3), and retinal pigment epithelial (RPE1-hTERT) cell lines (Fig. 2A). Specifically, we fixed cell lines two days after transfection and immunostained for ARL13B to identify which cells were ciliated. We then identified ciliated cells with visible plasmid-derived fluorescence and split them into three categories depending on whether fluorescence was exclusively localized to ARL13B-expressing primary cilia, localized to both primary cilia and other cellular compartments, or present in other cellular compartments but not detectable in primary cilia (Fig. S2B).

**Figure 2.**
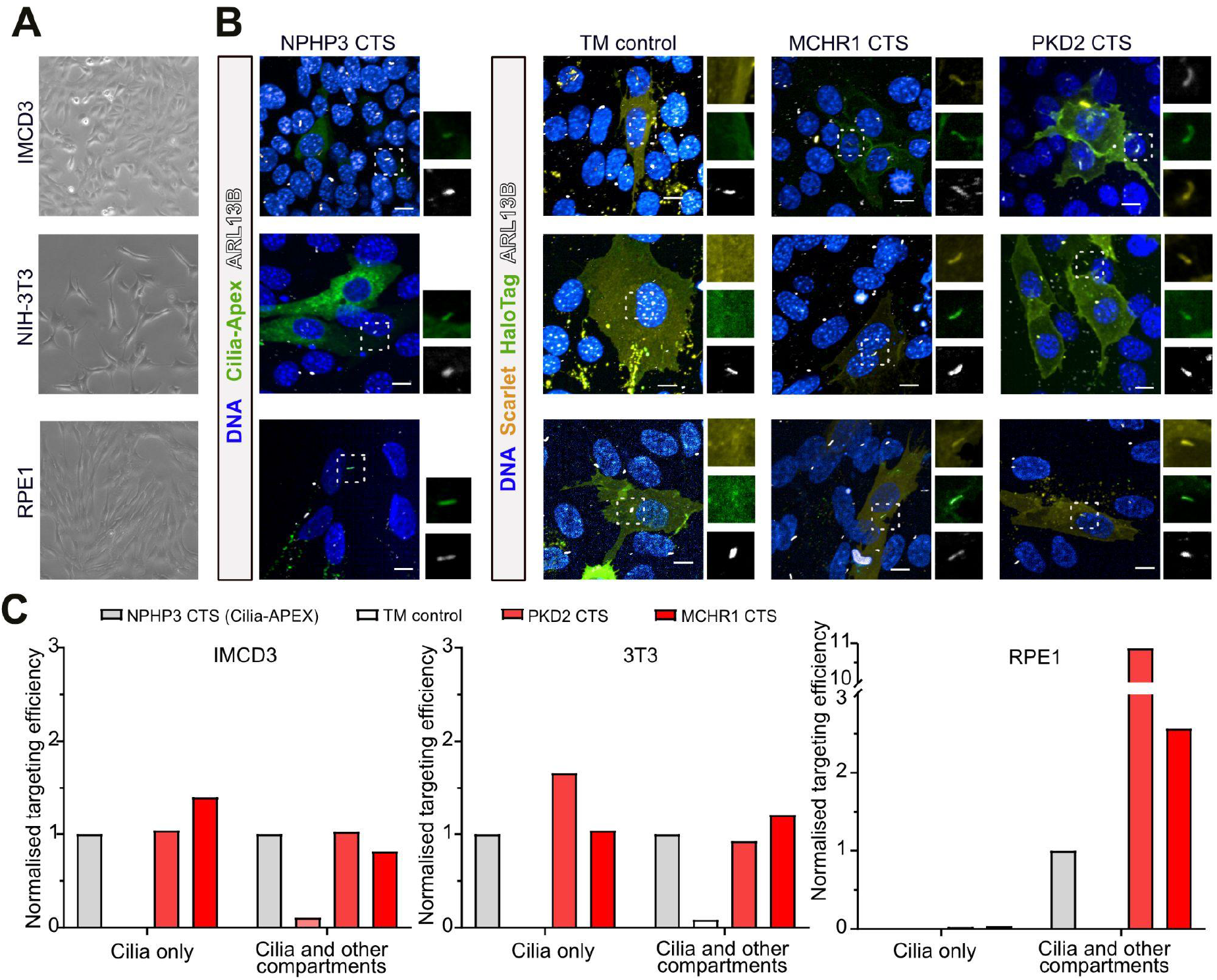
CTSs from MCR1 and PDK2 are sufficient for ciliary targeting in diverse cell lines in the context of a TMD. **A)** Phase contrast photomicrographs of ciliated IMCD3, NIH-3T3, or RPE cell lines used to test for ciliary localisation by transient transfection. **B)** Representative images of the subcellular localisation of experimental constructs as revealed by endogenous GFP fluorescence in cells transfected with NPHP3-containing Cilia-APEX construct, or mScarlet signal in cells transfected with control transmembrane (TM), or MCHR1- or PKD2-derived CTS-containing constructs. Scale bar is 10 µm for all images. **C)** Quantification of ciliary localisation for each tested cell line and constructs, normalized to the percentage of Cilia-APEX transfected cells where the fluorescent protein was exclusively observed in cilia, or observed in cilia and other cellular compartments, with the exception of the ‘cilia only’ category for RPE1 cells where no cells were identified for Cilia-APEX, n = 139 +/- 49 (IMCD3), 50 +/- 16 (3T3) and 81 +/- 41 (RPE1) averaged quantified cells per condition.

In agreement with previous reports, we found that Cilia-APEX containing the NPHP3 CTS localized to primary cilia in all tested cell lines, albeit at lower efficiency in RPE1 cells and less than 100% efficiency in all cell lines, likely due the variability associated with transient transfection (Fig. 2B, S2B). In parallel, we transfected the same cell lines with plasmids encoding our transmembrane constructs that either lacked a CTS (TM control), or contained MCHR1-3xCTS or PKD2-3xCTS. We then quantified the fraction of cells with exclusive or partial ciliary localisation as we had for Cilia-APEX, and found that MCHR1-3xCTS and PKD2-3xCTS performed at least as well (Fig. 2C, S2B), suggesting they are sufficient for ciliary targeting in a range of ciliated cell types, including human neurons.

### Generation of stably cilia-tagged human pluripotent stem cell lines

We hypothesized that the variable ciliary targeting efficiency we observed upon transient transfection could be improved by stably introducing a single copy of our ciliary targeting construct into a genetic ‘safe harbor’ locus. By targeting hiPSC lines, primary cilia could be labeled in any of their differentiated progeny, including cortical and hypothalamic neurons. We therefore extended the MCHR1-3xCTS transmembrane construct with a C-terminal ascorbate peroxidase (APEX2) to facilitate proximity labeling in future studies (Mick et al. 2015; May et al. 2021), and cloned it downstream of a CAG promoter into a targeting vector for the *AAVS1* safe harbor locus that facilitates transgene expression in hPSCs and their differentiated progeny (Hockemeyer et al. 2009) (Fig. 3A, S3A). We then introduced this construct, or a TM control construct lacking the MCHR1-3xCTS, into the deeply-characterized KOLF2.1J hiPSC line that differentiates well into neural cell types (Pantazis et al. 2022). We also targeted the *AAVS1* locus in a clone of the WTC-11 hiPSC line (Kreitzer et al. 2013) that had been previously modified to express neurogenin2 (*NGN2*) in a doxycycline-inducible manner to efficiently drive cortical-like neuronal differentiation (Fernandopulle et al. 2018). Upon drug selection, we picked clones, confirmed *AAVS1* targeting by PCR, and further modified targeted lines on the KOLF2.1J background by introducing a doxycycline-inducible *NGN2* expression construct by PiggyBac-mediated stable integration.

**Figure 3.**
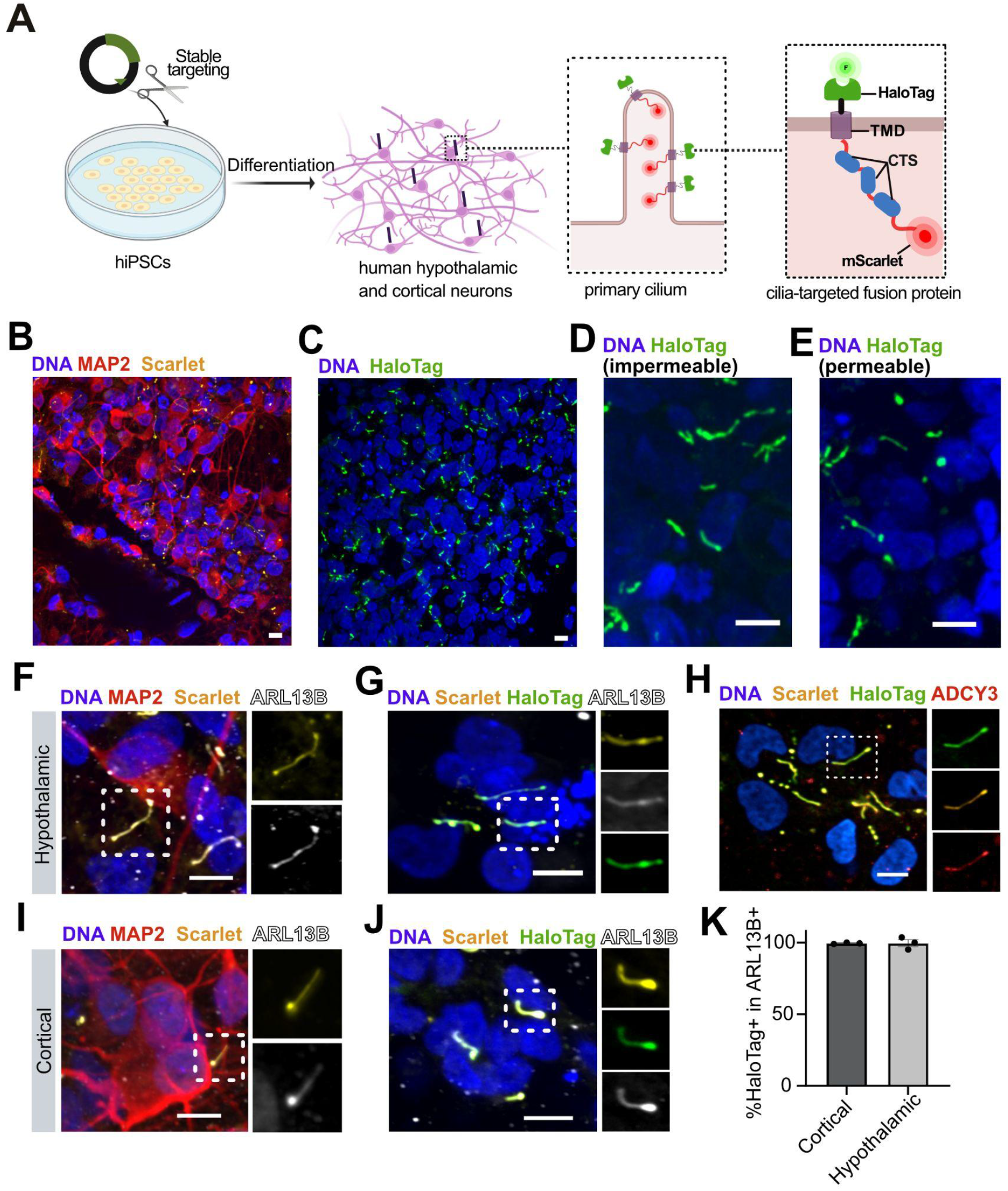
A stable hiPSC cell line constitutively expressing a primary cilia reporter. **A)** Experimental schematic for stable targeting of human *AAVS1* locus to introduce a constitutively expressed ciliary reporter. **B,C)** Low-magnification images of primary cilia in hiPSC-derived hypothalamic neurons visualized by mScarlet (B) or a fluorescent HaloTag ligand (C). **D,E)** Higher magnification images of cultures in (C) where cilia are visualized with cell-impermeable (D) or permeable (E) HaloTag ligands to confirm the specificity of the reporter signal. **F-H)** In hiPSC-derived MAP2+ hypothalamic neurons, ARL13B+ primary cilia can be visualized by both endogenous fluorescence of mScarlet (F), and fluorescent HaloTag ligand (G) in cilia that are immunopositive for ADCY3 (H). **I,J)** Similar results are seen in hiPSC-derived cortical neurons. Scale bars represent 10 µm in panels B-J. **K)** Fraction of ARL13B+ cilia that also are labeled by the HaloTag ligand in cortical and hypothalamic neurons. N = 3 technical replicates, n = 25 +/- 3 (hypothalamic) and 268 +/- 56 (cortical) ARL13B+ structures per replicate.

### Efficient ciliary targeting in stable cell lines

To assess the subcellular localisation of ciliary reporter constructs, we differentiated clones of KOLF2.1J and WTC-11 carrying both the stably-integrated *NGN2* transgene and either the TM or MCHR1-3xCTS into cortical neurons. After 10 days of differentiation, nearly all cells (93.4 +/- 4.6%) from both genetic backgrounds had adopted neuronal morphology and were immunopositive for the neuronal marker gene MAP2 (Fig. 3B). Upon imaging live cells, we found that the mScarlet signal did not localize to cilia in cells expressing the TM construct (Fig. S3B,C), but showed clear cilia-like localization in cells targeted with constructs containing MCHR1-3xCTS (Fig. 3B,C). Upon addition of fluorescent HaloTag ligands to live cells, we observed a bright and exclusively cilia-like fluorescence pattern for cells expressing the MCHR1-3xCTS (Fig. 3C), regardless of whether we added cell-permeable or cell impermeable ligands (Fig. D,E).

The expression pattern we observed appeared ciliary, but might only be labeling a subset of all cilia, or might be specific to cortical differentiation. We therefore differentiated cell lines to both cortical and hypothalamic neurons and fixed and immunostained cultures for the cilia-enriched markers ARL13B and/or ADCY3 (Fig. 3F-J). We then quantified the fraction of ARL13B immunopositive structures that were also positive for mScarlet and the Halo ligand (Fig. 3K), as well as the converse (mScarlet and HaloTag ligand-expressing structures that were also immunopositive for ARL13B, Fig. S3D). We observed nearly complete colocalization (>99%) of our construct with ARL13B+ primary cilia (Fig. 3H). We observed very similar results when we repeated this analysis in hiPSC differentiated to hypothalamic neurons (Fig. 3I-K), including ciliated POMC neurons (Fig. 3E). These results suggest that the cell lines we generated permit cilia to be studied and manipulated in a wide range of human cell types that can be generated from hiPSCs.

### Image-based screening for regulators of primary cilia length (PCL)

Since primary cilia can be readily visualized in live human neurons in our stably targeted cell lines, we reasoned they would be ideal for identifying factors that regulate primary cilia length (PCL). PCL dynamically changes in neurons and other cell types in response to a variety of exogenous signals that impact actin and microtubule polymerization and other pathways, though the precise mechanisms and functional consequences of PCL changes remain poorly understood (Macarelli, Leventea, and Merkle 2023). To test if PCL can be measured in hiPSC-derived cortical and hypothalamic neurons, we imaged live cultures labeled with fluorescent HaloTag ligand on the Opera Phenix automated confocal microscope. The bright and specific signal of this ligand in monolayers of adherent neurons allowed primary cilia to be readily detected and measured in an automated analysis pipeline (Fig. 4A). Specifically, we captured confocal stacks, took maximum intensity projections, and then automatically detected primary cilia and quantified their length (Fig. 4B). This automated pipeline enabled thousands of cilia (7850 +/- 434) to be measured in each well of a 96-well plate.

**Figure 4.**
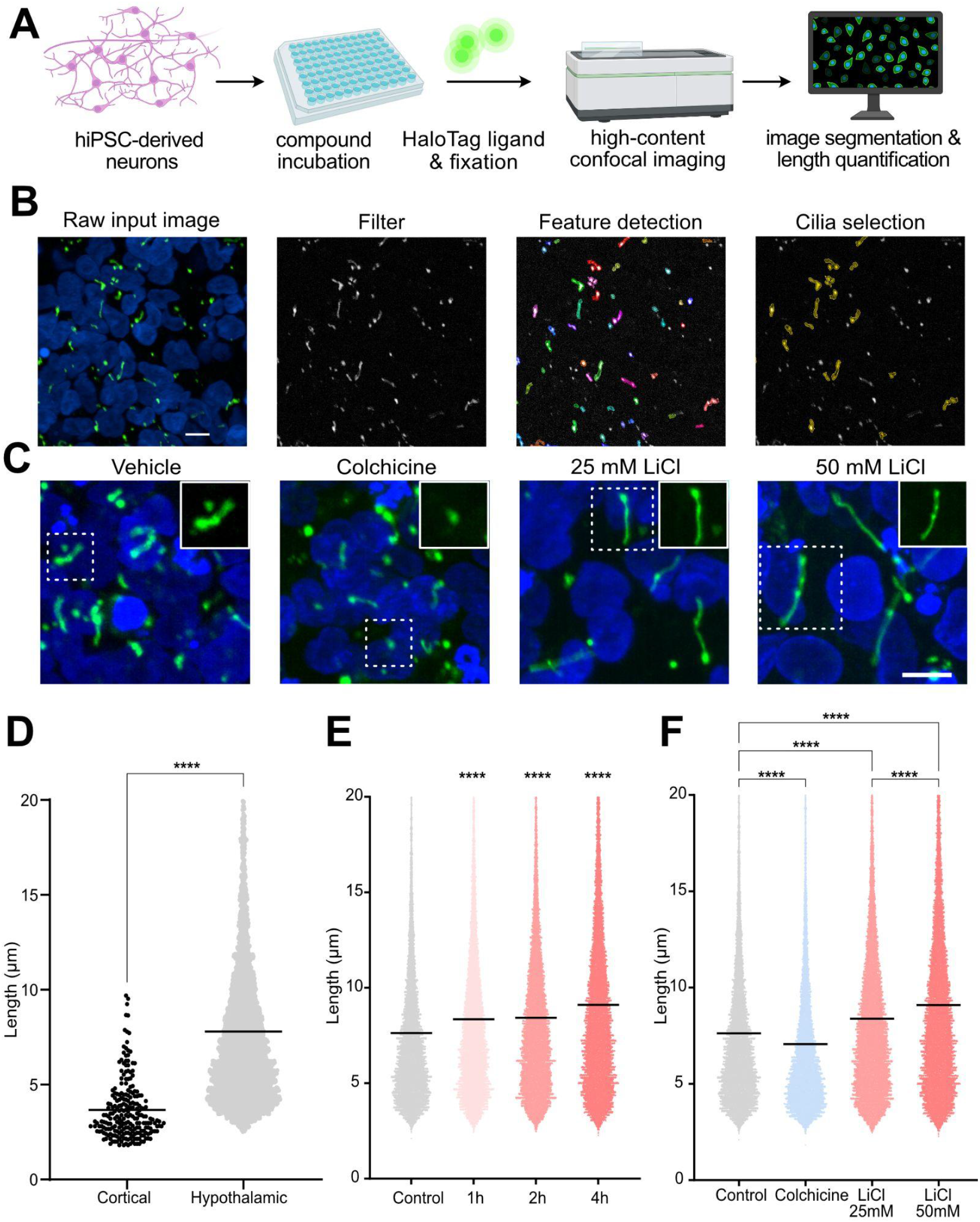
Modulation of primary cilia length (PCL) in human neurons. **A)** Experimental schematic for the automatic confocal image-based quantification of drug-induced changes in PCL in human neuronal cilia visualized by HaloTag ligand. **B)** Representative images of cell nuclei (blue) and fluorescent Halo ligand (green) that are filtered by thresholding prior to automatic segmentation and selection of objects that meet criteria expected of primary cilia, such as minimal area and length. **C)** Representative images from neuronal cultures treated with vehicle, 4 mg / mL colchicine, or 25 or 50 mM LiCl, where nuclei are in blue and the Halo ligand is in green. Scale bars represent 10 µm for B,C. **D)** PCL measurements in hiPSC-derived cortical and hypothalamic neurons, where each data point represents one primary cilium. Bar indicates mean cilia length. n = 235 and n = 5038 quantified structures/replicate for cortical and hypothalamic respectively. **E)** Time-dependent increase in measured PCL in response to 50 mM LiCl. **F)** PCL changes in response to treatment with vehicle, 4 mg/ml colchicine, or 25 or 50 mM LiCl for 4 hours. N = 3 technical replicates, 30 fields/replicate, and 7850 +/- 434 quantified structures per replicate. ****p<0.0001 by Kruskal-Wallis test.

To test whether this analysis pipeline can be used to measure changes in PCL, we treated hypothalamic neuron cultures with lithium chloride (LiCl), which is known to increase PCL in many cell types including neurons (Miyoshi et al. 2009), or with colchicine, an inhibitor of microtubule polymerisation that may destabilize cilia (Hierck et al. 2008; Rosenbaum and Carlson 1969). We observed apparent PCL changes (Fig. 4C) without apparent changes in primary cilia number per well (Fig. S4A). Since cultures of hypothalamic neurons had a higher fraction of ciliated cells and longer average cilia (7.64 +/- 0.14 µm) than cortical neurons in the conditions we tested (Fig. 4D), we proceeded to treat hypothalamic cultures with vehicle or 25 or 50 mM LiCl for 1, 2, or 4 hours, before adding fluorescent HaloTag ligand and fixing cells with 4% PFA (Fig. 4C, S4). We found that both 25 mM and 50 mM LiCl produced significant and time-dependent increases in PCL over the 4-hour treatment time course (Fig. 4E, S4B). Next, we selected the 4 hour time point and quantified changes in PCL in cultures treated with 4 mg/ml colchicine or LiCl and quantified the length of 90861 cilia. We found that primary cilia in wells treated with colchicine were significantly shorter than vehicle-treated controls (p<0.0001), whereas primary cilia in wells treated with 25 or 50 mM LiCl were significantly longer (p<0.0001) (Fig. 4F). Together, these results demonstrate the utility of the cilia reporter cell line for measuring PCL and pave the way for larger-scale studies to screen for factors that regulate PCL across diverse cell types.

## Discussion

We found that while commonly-used CTS failed in human neurons, we could identify two small CTSs that were sufficient to drive ciliary localisation of a fluorescent construct in every ciliated cell line we tested by transient transfection, and worked at efficiencies approaching 100% in stably targeted cell lines. Here, we discuss the potential uses of the stable hiPSC cell line to study the biology of primary cilia, the reasons these sequences were effective in neurons while others were not, and the strengths and limitations of our study.

The construct we stably targeted to hiPSCs permits live imaging of cilia via mScarlet fluorescence or fluorescent Halo ligands, pull-down or other manipulation of cilia via the extracellular Halo tag, and proximity proteomics of primary cilia. We also demonstrate that one can test for factors that alter primary cilia length by confirming the known lengthening effects of LiCl, and showing that colchicine treatment is sufficient to reduce primary cilia length under these conditions. The bright and specific ciliary staining obtained by the HaloTag ligand improves upon cilia identification by immunostaining in both its high signal to noise ratio, and its ability to be applied to live cells rather than fixed cultures. These and other analyses made possible by stable ciliary reporter cell lines could be extended to any cell type that can be generated from hiPSCs.

In our hands, a Cilia-Apex construct containing the myristoylation site and coiled-coil domains of NPHP3, previously reported to mediate ciliary targeting, (Nakata et al. 2012) did not show this activity in iPSC-derived human neurons. We speculate that there are cell type-specific mechanisms that are either less efficient or absent in the neurons we tested. The similar features of the two successful constructs containing CTSs derived from MCHR1 and PKD2 sheds some light on these targeting mechanisms, as both contain a CTSs also derived from transmembrane proteins, suggesting both are required for membrane protein sorting to the primary cilium. Previous studies suggest that BBS proteins may be important for MCHR1 ciliary targeting, since this receptor accumulates in cytoplasmic puncta in *Bbs2* and *Bbs4* double knockout mutant mice (Berbari, Lewis, et al. 2008). Future immunoprecipitation experiments with the CTSs from this study could reveal shared interaction partners.

It is not clear why other tested CTS did not work in our system. The sequence from KIF17 may have been insufficient when tested without the kinesin motor that appears to promote its association with importin-β2 and transport across the ciliary transition zone (Dishinger et al. 2010). The CTS in NPHP3 and CYS1 both require myristoylation, and although these sequences were included in the sites we tested, it is possible that this post-translational modification did not proceed efficiently in iPSC-derived neurons. RHO requires a complex of TNPO1 and RAB8 (Madugula and Lu 2016; Follit et al. 2010) and/or dual Ax(S/A)xQ motifs for ciliary targeting, and the human RHO CTS tested contains only one of these motifs (Geneva, Tan, and Calvert 2017). NPY2R and SSTR3 both localize to neuronal primary cilia, but the conformation of the isolated CTS might be different than in the native protein, additional sequence context might be necessary for the isolated CTSs to engage with the ciliary targeting machinery, or perhaps post-translational modifications in the CTS differ between the native protein our artificial construct.

## Limitations of study

We could not exhaustively test all potential CTSs, and cannot exclude the possibility that more potent or universally effective CTS remains to be discovered. To increase our chances of success, we tested CTS in a 3x array of putative CTS with a ‘core’ sequence important for ciliary targeting, and flanked with surrounding residues to provide sequence context. Future studies could test whether a single copy of the CTS is sufficient to mediate ciliary targeting, or systematically mutate or delete residues in the tested CTS to identify those that are necessary for the observed targeting. These studies might enable more compact or effective CTS to be designed and provide insights into the mechanisms of ciliary targeting.

Of the constructs we tested that included a TMD, the candidate CTSs were separated from the membrane by a short sequence (KRKRGS). As the TMD was required for ciliary targeting, it is possible that the artificial nature of our TMD (string of leucine residues), the positively charged residues following this sequence, or the proximity of the CTS to the membrane contributed to the observed activity. Similarly, we did not test constructs targeted to the membrane by lipid modification, which could engage distinct mechanisms of ciliary localization. The IL2 signal peptide is widely used to target proteins to the secretory or transmembrane pathways (Zhang, Leng, and Mixson 2005), and our staining with cell-permeable and cell-impermeable Halo ligands suggests that our construct efficiently inserted into the plasma membrane the HaloTag facing the outside of the cell. Systematically testing other signal peptides might shed light onto their cooperative or antagonistic interactions with cytoplasmic CTS.

Although we tested two different methods of generating neurons from iPSCs and obtained similar results on two different genetic backgrounds, it is possible that some of the effects we observed could be due to the relatively immature state of cells derived from iPSCs. Finally, when generating stable cell lines, we targeted a single copy of the construct to the *AAVS1* locus under the control of the strong and constitutive CAG promoter, and acknowledge that its presence at the cilium might alter endogenous protein localisation, either by physically crowding out ciliary proteins or competing for access to the ciliary localisation and transport machinery.

In summary, the versatility of the short CTS we describe should enable groups to target constructs of their interest to primary cilia in their preferred cell types and model systems. The short size of the CTS we describe makes it an attractive candidate for generating fusion proteins that must remain compact due to steric limitations or viral genomes for use in animal models. As a resource to the community, the constructs, cell lines and analysis pipelines described in this study are freely available upon request.

## Materials and methods

### Cell culture

All cultured cells in this study were maintained at 37°C in humidified incubators at 5% CO_2_ and 20% O_2_. Human induced pluripotent stem cell (hiPSC) lines KOLF2.1J (P24 to P38) (Pantazis et al. 2022) and WTC-11 (P40 to P45) (Kreitzer et al. 2013) were maintained as previously described (H.-J. C. Chen et al. 2023). Briefly, cells were maintained in StemFlex medium on plates coated with Geltrex, and passaged with 0.5 mM EDTA (Ethylenediaminetetraacetic acid) when 70-80% confluent, and replated in the presence of 10 µg/mL Rock Inhibitor Y-27632. The IMCD3 cell line (P20-30) (Mukhopadhyay et al. 2010) was maintained in DMEM-F12 medium supplemented with 2.5 mM L-Glutamine, 10% fetal bovine serum (FBS) and 1% Penicillin-Streptomycin. NIH-3T3 (3T3) fibroblasts (P5 to P10) were maintained in DMEM medium supplemented with 10% FBS, while RPE1-hTERT cells (P7 to P12) were maintained in DMEM-F12 medium supplemented with 2.5 mM L-Glutamine, 10% FBS and 1% Penicillin-Streptomycin. IMCD3, 3T3 and RPE1 cells were passaged with Trypsin - EDTA solution when 70-80% confluent.

### Cortical neuron differentiation

Cortical neurons were generated by inducibly expressing the transcription factor neurogenin-2 (NGN2) with doxycycline as previously described (Fernandopulle et al. 2018) and detailed below. The WTC-11-NGN2-dCAS9 hiPSC line, in which NGN2 transgene was stably integrated into the *AAVS1* safe-harbor locus under a doxycycline-inducible promoter and dCAS9 was inserted into the *CLYBL* safe-harbor locus (Tian et al. 2019), was a gift from Michael E. Ward. The KOLF2.1J-NGN2 hiPSC line was generated by two consecutive rounds of transfection with a piggyBac transposon vector containing human *NGN2* under a dox-inducible promoter (Laraflores et al., 2022) together with a plasmid encoding the piggyBac transposase in a 3:1 ratio (vector:transposase) using the P3 Primary cell 4D-Nucleofector™ kit (Lonza). Prior to targeting, all plasmid DNA was purified by Triton X-114 isothermal extraction as described previously (Ma et al. 2012). A total of 3 µg DNA was delivered to 8x10^5^ hiPSCs dissociated to a single-cell suspension with TrypLE Express, and cell were plated into one well of a 6-well plate in the presence of 10 µg/mL Rock Inhibitor Y-27632. After allowing the cells to recover for 72 hours, they were exposed to 5 μg/mL of blasticidin for six days to select for cells that had stably integrated the NGN2 transgene. Finally, twelve clonal lines were isolated, validated by assessing the efficiency of neuronal differentiation and one was selected to use for experiments.

Human iPSCs were dissociated with 1X TrypLE Express and differentiated for three days in neuronal induction medium (DMEM/F12 with HEPES, N-2 Supplement, Glutamax, MEM non-essential amino acids (NEAA) supplemented with 2 µg/mL Doxycycline, 10 µg/mL Rock Inhibitor Y-27632, and 50 µM 2-Mercaptoethanol) at a density of 1x10^5^ cells per cm^2^ on Geltrex-coated plates. Rock Inhibitor was withdrawn after 24h. After three days, cells had adopted neuronal morphology and were dissociated with a mixture of 1x TrypLE Express and 200 U/mL papain (10:1) and washed with medium supplemented with 2 mg/mL DNAse I (1:60), and replated in cortical medium at 3x10^5^ cells per cm^2^ onto plates coated with 0.1 mg/mL Poly-Ornithine. Cortical medium consisted of BrainPhys medium containing B27 Supplement, 10 ng/mL human neurotrophic factors (hGDNF, hBDNF and hNT-3), 1 µg/mL mouse Laminin and 10 µg/mL Rock Inhibitor Y-27632. For neuronal maintenance, 50% cortical medium was replaced every 2-3 days until day 10 to 20 of differentiation, when neurons were used for experiments. When differentiating the KOLF2.1J PiggyBac line, 1 µM uridine and 1 µM fluorodeoxyuridine were added to the media from day 2 onward to suppress mitotic cells.

### Hypothalamic neuron differentiation

Human iPSCs were differentiated to hypothalamic neurons by directed differentiation as previously described (H.-J. C. Chen et al. 2023; Merkle et al. 2015; Kirwan, Jura, and Merkle 2017). Briefly, hiPSCs were dissociated with TrypLE Express and plated onto Geltrex-coated plates at a density of 1x10^5^ cells/cm^2^ in StemFlex medium supplemented with 10 µg/mL Rock Inhibitor Y-27632. The following day, medium was changed to N2B27 medium (1:1 Neurobasal-A to DMEM/F12 with GlutaMAX media, 1X Glutamax, 0.075% Sodium bicarbonate, 1X MEM NEAA, 200 mM ascorbic acid, 1X penicillin-streptomycin, 1X B27 supplement and 1x N2 supplement). Small molecule modulators of developmentally important signaling pathways were then sequentially added to induce hypothalamic fate as previously described (H.-J. C. Chen et al. 2023). On day 13 of differentiation, hypothalamic progenitors were dissociated with a mixture of 1X TrypLE Express and 200 U/mL papain (10:1), washed with medium supplemented with 2 mg/mL DNAse I (1:60), and re-plated at a concentration of 3x10^5^ cells/cm^2^ onto 4 μg/mL Laminin 511-coated plates in N2B27 medium (see above) supplemented with 10 μg/mL BDNF and 10 µg/mL Rock Inhibitor Y-27632. The following day, media was changed to Synaptojuice medium 1 (N2B27 medium, 2 μM PD0332991 isethionate, 5 μM DAPT, 370 μM CaCl_2_ , 1 μM LM22A4, 2 μM CHIR99021, 300 μM GABA, and 10 μM NKH447) supplemented with 10 μg/mL BDNF. After 1 week, media was switched to Synaptojuice 2 (N2B27 medium, 2 μM PD0332991 isethionate, 370 μM CaCl_2_, 1 μM LM22A4, and 2 μM CHIR99021) and cells were cultured for at least another week. Complete medium change was performed every 2 days and cells were used for experiments from day 30 of differentiation onwards.

### Cloning of fusion protein constructs containing candidate CTSs

To test the ability of candidate ciliary targeting sequences (CTSs) to localize a fluorescent protein (mScarlet-I) to primary cilia, DNA fragments containing three repetitions of a CTS from known ciliary proteins (Melanin-concentrating hormone receptor 1 (MCHR1), Polycystin 2 (PC2), Cystin (CYS1), Somatostatin receptor 3 (SSTR3), Kinesin family member 17 (KIF17), Neuropeptide Y receptor Y2 (NPY2R) and Rhodopsin (RHO) flanked by flexible glycine and serine linkers (GGGGS) were synthesized as double-stranded DNA by Integrated DNA Technologies and Twist Bioscience. Each fragment was then cloned into the pScarlet-N1 plasmid (RRID:Addgene_128060) from Oskar Laur, previously linearized by digestion with BmtI and PstI, using the In-Fusion HD EcoDry Cloning Plus kit (Cyt-Cilia constructs). The resulting plasmids were verified by restriction digestion and Sanger sequencing, and we referred to those with expected sequences as ‘cytoplasmic’ constructs. The unmodified mScarlet vector served as a negative control.

To generate fusion proteins able to localize to the ciliary membrane, the Interleukin-2 (IL-2) signal peptide, a HaloTag extracellular domain and a transmembrane domain (TM) were synthesized ahead of the 3xCTS array (see above) as double-stranded DNA by Twist Bioscience. The TM domain consisted of 22 Leucine amino acids followed by four positively-charged amino acids to stabilize the TM domain in the plasma membrane and aid export through the biosynthetic secretory pathway (H. Chen and Kendall 1995). A DNA fragment lacking only the CTSs array was also synthesized as a control (TM control). Each fragment was then cloned into the pScarlet-N1 plasmid as described above, and resulting plasmids were verified by restriction digestion and Sanger sequencing. We referred to those with expected sequences as ‘transmembrane’ constructs. A plasmid containing the IL2 signal peptide, HaloTag, TM domain, and mScarlet but lacking a CTS served as a negative ‘TM’ control.

### Transient transfection in cell lines and hPSC-derived neurons

To test for the ciliary localization of the candidate plasmids described above or control plasmids including Cilia-APEX, all plasmid DNA was purified by Triton X-114 isothermal extraction as described previously (Ma et al. 2012) to remove endotoxin and transiently transfected into different cell types. Cilia-APEX (pEF5B-FRT-cilia-APEX) enables GFP and APEX to be targeted to cilia in many cell types via the N-terminal 203 residues of NPHP3 and was a gift from Maxence Nachury (RRID:Addgene_73186) (Mick et al. 2015). IMCD3, 3T3 and RPE1 cells were transiently transfected with these and our cloned constructs using Lipofectamine™ LTX Reagent according to manufacture’s instructions when 70-80% confluent. Prior to transfection, the FBS in the media was reduced to 0.2% and cells were then serum-starved for the following 48h. Cells were then fixed and imaged. Due to the low efficiency of lipofection in human neurons, plasmids were introduced to these cells by electroporation. Specifically, 1x10^6^ hiPSC-derived neurons were dissociated with 1x TrypLE Express supplemented with 200 U/mL papain (10:1) and resuspended in 100 μl 1x Opti-MEM I Reduced Serum medium with 2 μg endotoxin-free plasmid DNA. Electroporation was performed using the NEPA21 electroporator (Poring Pulse: 150 V, 5 ms length, 50 ms interval, 2 times; Transfer Pulse: 20 V, 50 ms length, 50 ms interval, 5 times). After electroporation, cells were resuspended in 400 μl of neuronal maturation medium supplemented with 10 μM Rock inhibitor, and plated at density of 6x10^5^ cells per cm^2^ in 96-well PhenoPlate microplates. Half medium was changed daily for 72 hours, until cells were fixed and imaged.

### Generation of stable hPSC lines carrying the ciliary targeting construct

To generatie targeting vectors for stable cell line generation, we performed site-directed mutagenesis on pEF5B-FRT-cilia-APEX (RRID:Addgene_73186) to convert APEX to APEX2, and then PCR amplified the coding sequence of APEX2 and cloned it into the Age1 restriction site downstream of mScarlet to produce TM-Scarlet-APEX2, TM-PKD2(3xCTS)-Scarlet-APEX2, and TM-MCHR1(3xCTS)-Scarlet-APEX2 plasmids. These sequences were then PCR amplified, and PCR products were digested with DpnI and cloned into the pENTR™ TOPO^R^ Vector using the pENTR™/D-TOPO™ Cloning Kit. The resulting pENTR plasmids were then used to efficiently shuttle the constructs into a destination vector using Gateway™ LR Clonase™ II to generate the plasmids to target a single copy of each construct to the adeno-associated virus integration, intron 1 (*AAVS1*) locus of hiSPC lines.

The *AAVS1* destination vector was assembled by PCR amplification and Gibson assembly from the pDTA-TK from Robert Benezra’s group (RRID:Addgene_22677) (Nam and Benezra 2009) to introduce negative selection cassettes flanking the *AAVS1* homology arms from the Gateway-compatible pAAVS1-P-CAG-DEST vector from Knut Woltjen (RRID:Addgene_80490). The resulting vector (pAAVS1-CAG-DEST-DTA-TK) enables constructs cloned into the attR sites to be driven by the strong, constitutive CAG (CAGGS) promoter (Oceguera-Yanez et al. 2016).

Prior to targeting hiPSCs, all plasmid DNA was purified by Triton X-114 isothermal extraction as described previously (Ma et al. 2012). Cell lines were then targeted at the *AAVS1* locus to generate AAVS1-TM-MCHR1(3xCTS)-mScarlet-APEX2 and AAVS1-TM-mScarlet-APEX2 cell lines using zinc finger nuclease (ZFN) mediated genome editing, as described previously (Chambers et al. 2009). Briefly, KOLF2.1J and WTC-11 hIPSC were electroporated using the P3 Primary Cell 4D-Nucleofector kit at a 2:1 ratio of *AAVS1* targeting vector:ZNF DNA per reaction. A plasmid encoding ZFNs targeting the *AAVS1* locus (pCMV-ZFN-AAVS1) was a gift from Dmitriy Mazurov (RRID:Addgene_89707) (Zotova et al. 2017). KOLF2.1J was targeted with both constructs, whereas the WTC-11 line was targeted only with the AAVS1-TM-MCHR1(3xCTS)-mScarlet-APEX2 construct. After nucleofection, cells were plated onto plates coated with 25 µg/ml Synthemax in StemFlex media supplemented with 10 µg/ml Rock Inhibitor Y-27632 and 1 μM Alt-R™ HDR (homology directed repair) Enhancer V2 and cultured in a 32 °C/5% CO2 incubator to cold shock cells and enhance HDR efficiency (Skarnes, Pellegrino, and McDonough 2019). The media was then replaced with StemFlex media without Y-27632 or HDR enhancer the next day (24h) and the cells were moved back to the 37 °C/5% CO2 incubator after 3 days (72h). When cultures reached approximately 70% confluence, media was changed daily for five days with StemFlex supplemented with 0.5 μg/ml puromycin dihydrochloride to select for stably modified cell clones. After allowing resulting single-cell derived clones to expand, clones were picked into 96 well plates, expanded, and tested for stable integration at the *AAVS1* locus by PCR amplification from genomic DNA as previously described (Santos et al. 2016).

### Immunofluorescence and imaging

To visualize the HaloTag in live cells, cultures were incubated with 200 nM Janelia Fluor HaloTag Ligand (Grimm et al. 2015) JF503/635i or 1μM Promega HaloTag Alexa Fluor 488 Ligand for 15 minutes at 37°C. Cells were washed 2 times with media and 1 time with PBS, then fixed as described below. For immunofluorescence, neurons derived from hiPSCs, and IMCD3, 3T3 and RPE1 cell lines were cultured on PhenoPlate 96-well microplates, fixed in 4% paraformaldehyde for 20 minutes at room temperature, and washed once with phosphate-buffered saline (PBS). IMCD3, 3T3 and RPE1 cell lines were then permeabilized with cold methanol at -20°C for 10 minutes and rinsed three times with PBS. Cells were then incubated overnight at 4°C with primary antibodies diluted in a blocking solution composed of PBS with 0.1% Triton X-100 (PBST) and 1% sterile-filtered normal donkey serum (NDS). Cells were then washed three times with PBS and incubated with secondary antibodies diluted in blocking solution for 2 hours at room temperature. Finally, cells were washed three times with PBS and incubated for 5 minutes at room temperature with 300 μM of 4′,6-diamidino-2-phenylindole (DAPI) in PBS before being washed with PBS and immediately imaged, or stored in PBS with 0.1% sodium azide.

Images were acquired with the Perkin Elmer Opera Phenix Plus High-Content Screening System with a 40x water objective, with 10-15 optical planes per Z-stack at 1 μm intervals. Images were processed with the ImageJ software to create single maximum projection image Some photomicrographs were background-subtractedto aid in visualization without adjusting gamma values. Some figure elements were prepared with the help of BioRender.com, and figures were assembled using Affinity Designer software.

### Quantification of ciliary labeling efficiency

Ciliary targeting efficiency in IMCD3, 3T3 and RPE1 cells was assessed in all cells that had detectable fluorescence from the transfected construct, had primary cilium immunopositive for ARL13B, and were located completely within imaging field borders. Among these cells, we defined ‘cilia targeting’ as the percentage of cells in which mScarlet fluorescence was detected in either just the ARL13B+ cilium, the ARL13B+ cilium and other cellular compartments, or only other cellular compartments. In hiPSC-derived neurons stably targeted with the cilia reporter construct (or transmembrane control constructs), we immunostained cultures for ARL13B (488 or 647 nm), counterstained with DAPI (405 nm), exposed cells to HaloTag ligand (488 or 635 nm), and imaged these channels and endogenous mScarlet (555 nm) signal. We then quantified the fraction of ARL13B+ cilia that co-expressed either the HaloTag ligand or mScarlet, and also quantified the fraction of HaloTag ligand positive or mScarlet positive ciliary structures that were immunopositive for ARL13B.

### Cilia length quantification

Primary cilia length in hiPSC-derived neurons stained with DAPI (405 nm) and HaloTag ligand (488 nm) was measured using the Harmony High-Content Imaging and Analysis software (Perkin Elmer, v4.9) using an automated analysis pipeline. Briefly, a fixed-intensity threshold was applied to every image in the 488 (HaloTag ligand) channel to exclude background noise, followed by the automated detection of objects with an area bigger than 4 µm^2^, roundness value lower than 0.8, and intensity value higher than 36. Objects falling on imaging field borders were excluded from analysis. Finally, cilia number and the length of HaloTag ligand+ objects (in microns) was measured. Detailed analysis workflows are available upon request.

### Statistical analysis

GraphPad Prism (Version 10.2.0) was used for statistical analysis. Data are presented as mean +/- standard error of the mean (SEM) or individual values +/- SEM. ANOVA followed by Bonferroni post hoc test or Kruskal–Wallis test followed by Dunn’s post hoc test were used to compare more than two datasets. Mann-Whitney test was used to compare two datasets. P values < 0.05 were considered to be statistically significant.

#### Key resources table

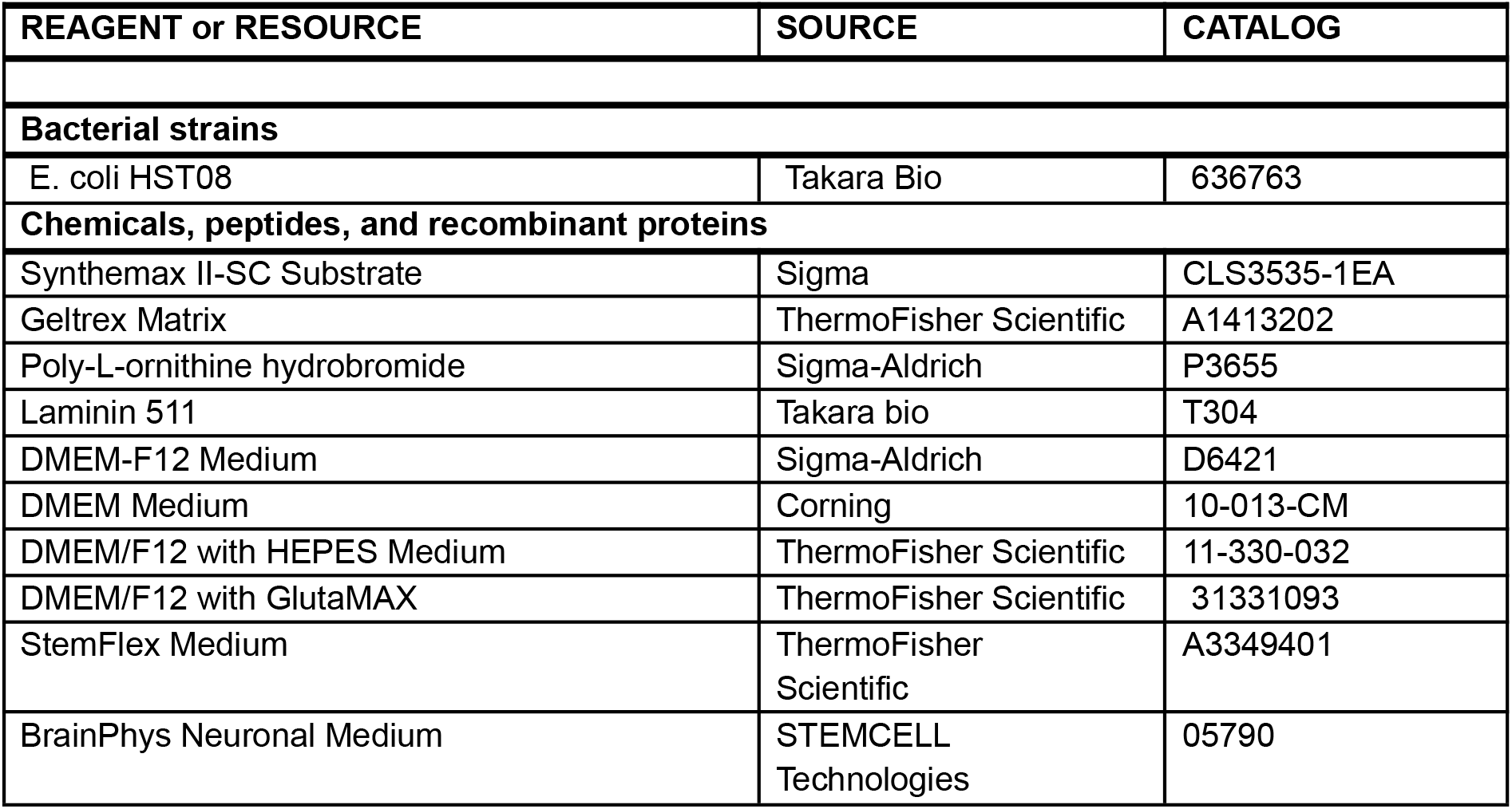
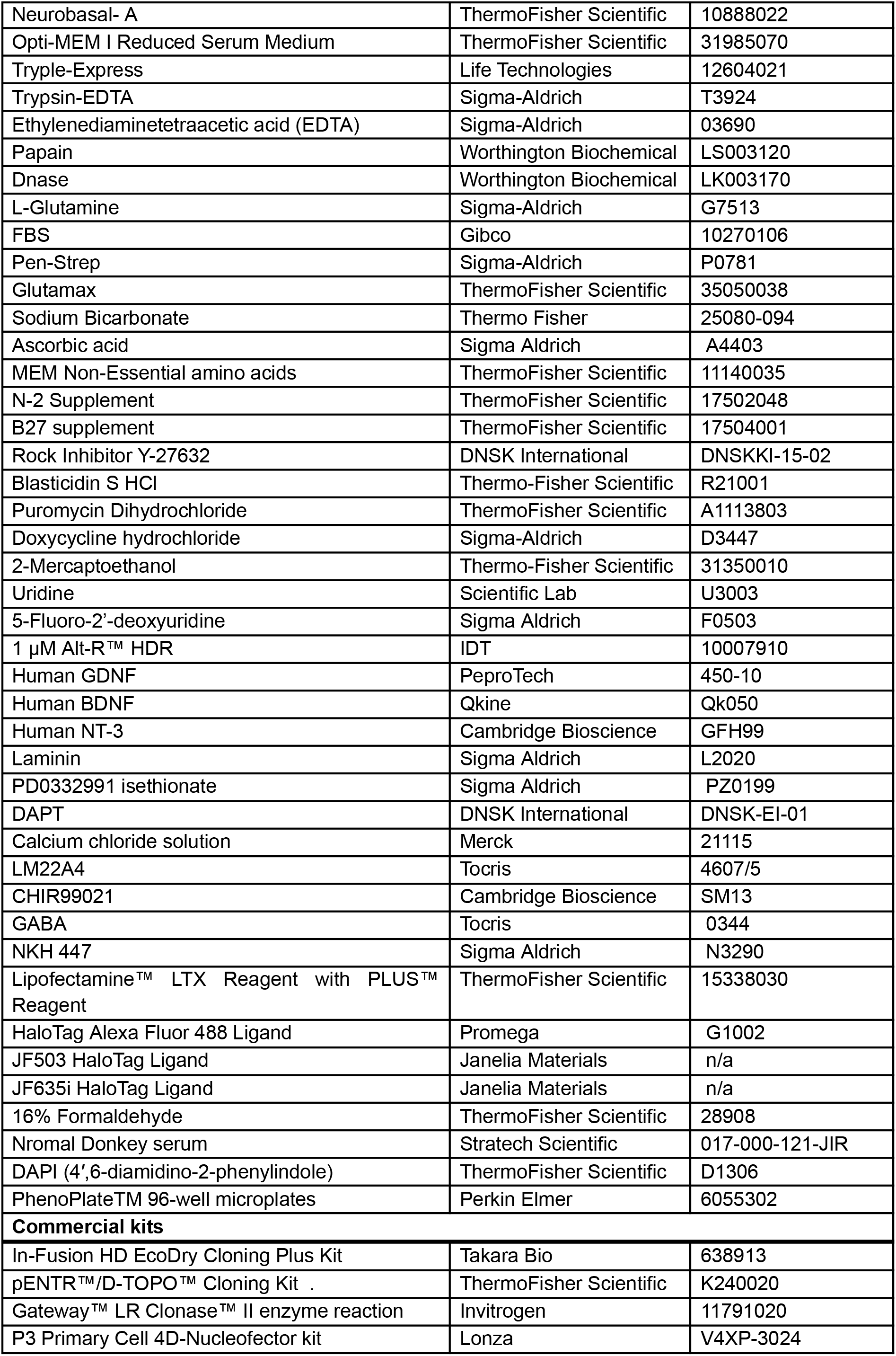
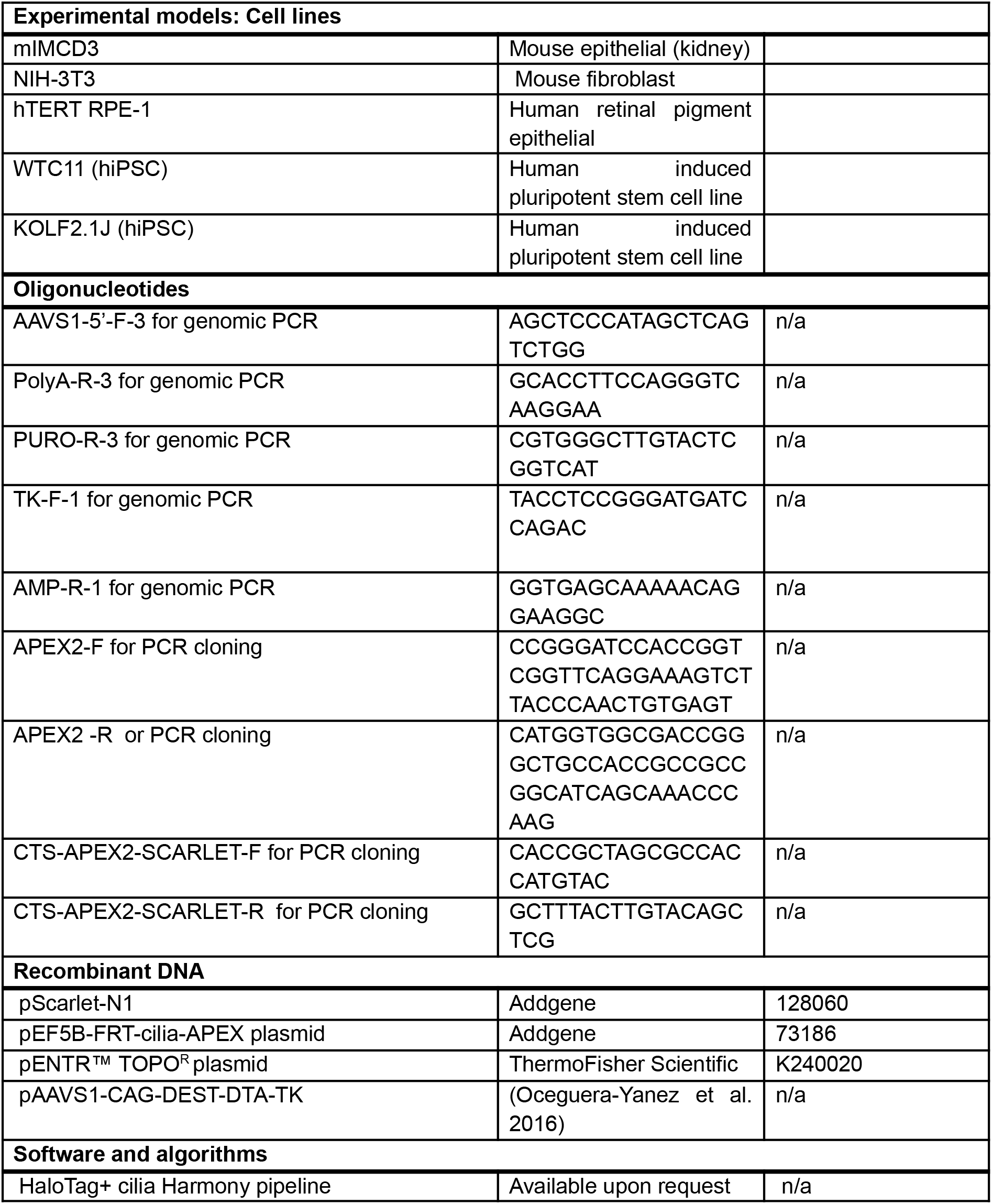

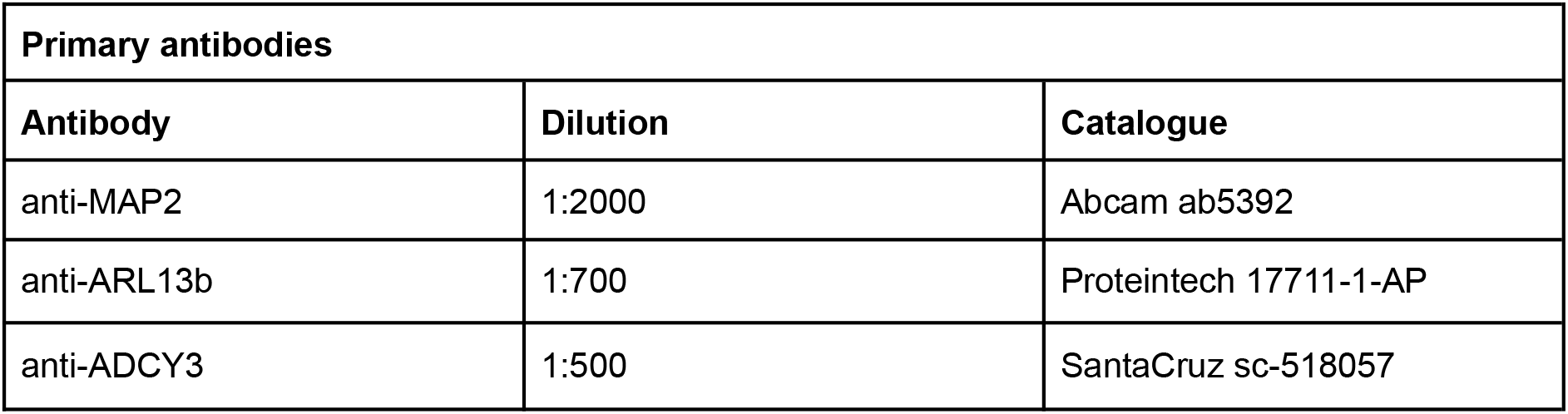
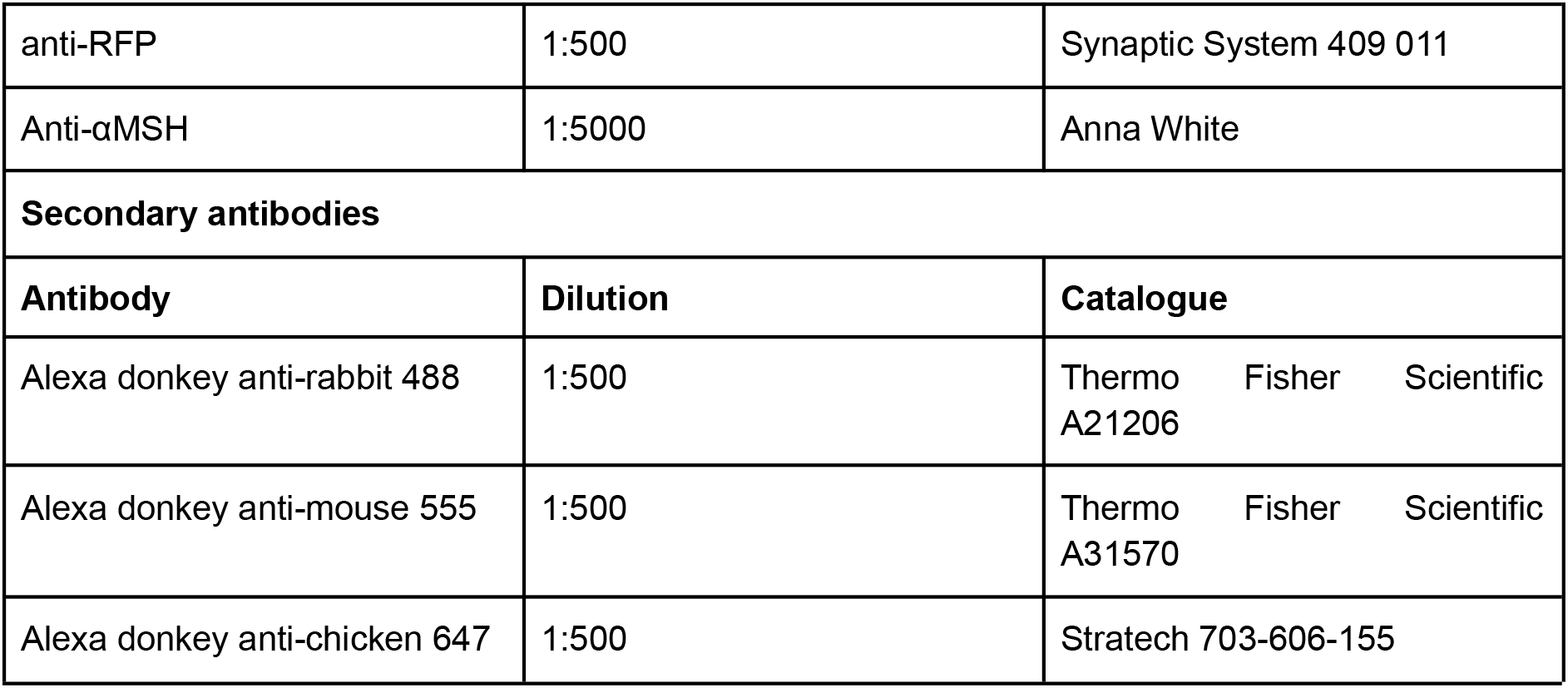

| **Primary antibodies** |  |  |
| --- | --- | --- |
| **Antibody** | **Dilution** | **Catalogue** |
| anti-MAP2 | 1:2000 | Abcam ab5392 |
| anti-ARL13b | 1:700 | Proteintech 17711-1-AP |
| anti-ADCY3 | 1:500 | SantaCruz sc-518057 |
| anti-RFP | 1:500 | Synaptic System 409 011 |
| Anti-αMSH | 1:5000 | Anna White |
| **Secondary antibodies** |  |  |
| **Antibody** | **Dilution** | **Catalogue** |
| Alexa donkey anti-rabbit 488 | 1:500 | Thermo Fisher Scientific A21206 |
| Alexa donkey anti-mouse 555 | 1:500 | Thermo Fisher Scientific A31570 |
| Alexa donkey anti-chicken 647 | 1:500 | Strattech 703-606-155 |

## Acknowledgements.

We thank David Mick for generously providing the Cilia-Apex construct, the IMCD3 FLP-In line, and helpful discussions that improved this manuscript. We thank Anne White for providing anti-αMSH antibody, Michael Ward, Mariam Aslanyan, Ronald Roepman, and Jiami Guo for sharing reagents, Ilona Zvetkova for her help in the design of the cloning strategies, Antony Hynes-Allen and Eugene Seah for their assistance with cloning, and Gregory Strachan for his initial assistance with the Opera Phenix confocal microscope and Harmony analysis software. F.T.M. is a New York Stem Cell Foundation - Robertson Investigator [NYSCF-R-156] and Chan Zuckerberg Initiative - Ben Barres Early Career Investigator [CZI NDCN 191942, 10.37921/429861umrcjh], and is supported by the Wellcome Trust and Royal Society [211221/Z/18/Z]. D.C.G. is funded by a Sir Henry Dale Fellowship from the Wellcome Trust/Royal Society (Grant 210481) and a Biotechnology and Biological Sciences Research Council responsive mode grant (BB/W005905/1). V.M. is supported by the Sackler Trust, the Medical Research Council Doctoral Training Programme and the School of Clinical Medicine Cambridge Trust Scholarship. For the purpose of open access, the authors have applied a CC-BY public copyright license to any Author Accepted Manuscript version arising from this submission.

## Author contributions

Conceptualization, Methodology, Investigation, Validation, Writing (original draft): V.M., and F.T.M. Funding Acquisition, Supervision, and Project Administration: F.T.M. Generation of Resources, V.M. and D.G. Formal Data Analysis and Visualisation: V.M., E.H., and F.T.M. Writing (Reviewing & Editing): all co-authors.

## Inclusion and diversity

We support inclusive, diverse, and equitable conduct of research.

---

**Figure S1.**
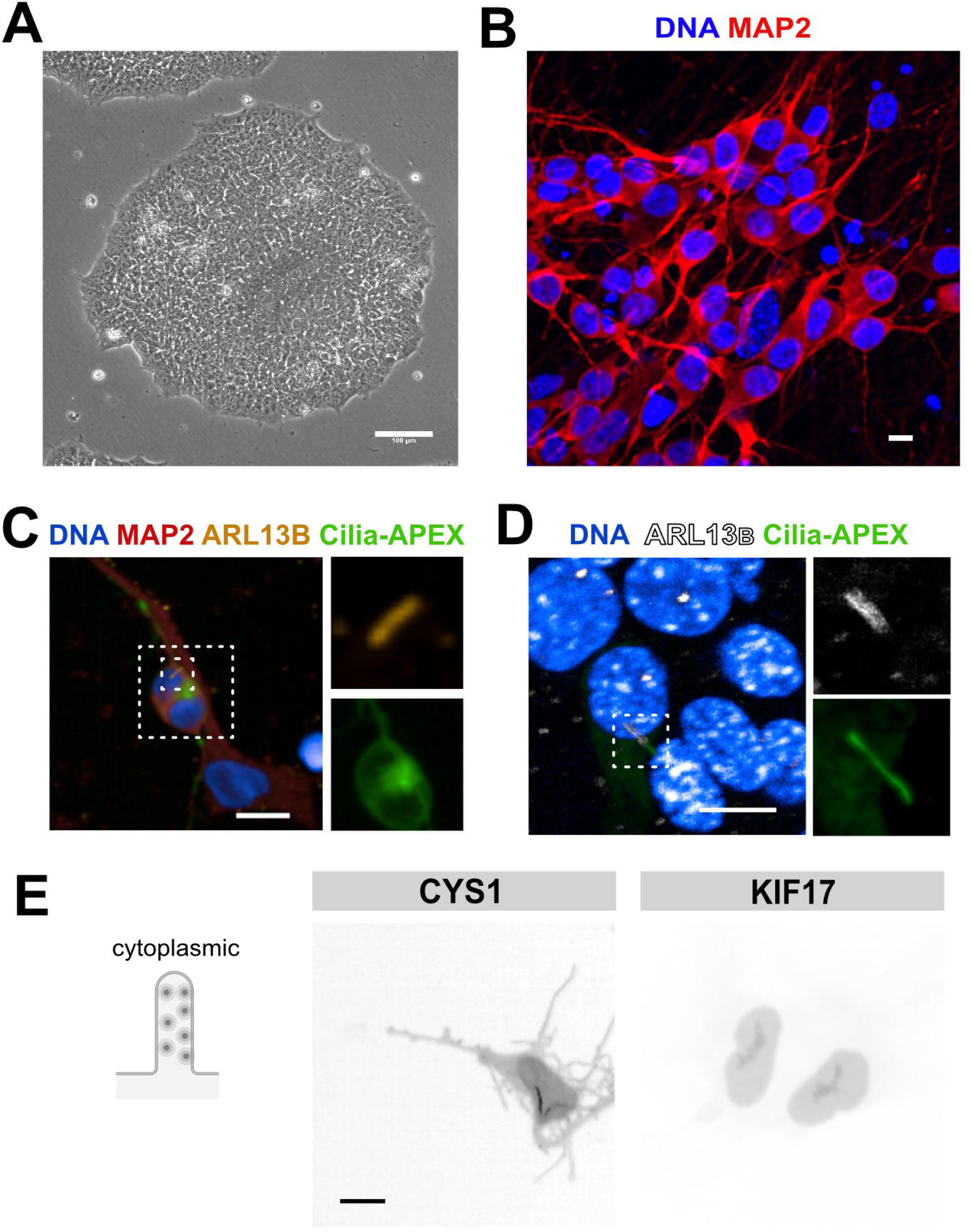
Neuronal differentiation of iPSCs and testing of ciliary localisation. **A)** Representative image of a colony of KOLF2.1J hiPSCs. **B)** Human iPSC-derived neurons immunostained for the marker MAP2. **C,D)** Upon transient transfection of Cilia-APEX into neurons, GFP signal is seen throughout the cell but not selectively localized to primary cilia (C), but the construct localizes exclusively to cilia in IMCD3 ciliated cell line (D). **E)** Cells transfected with cytoplasmic constructs with CTSs derived from CYS1 and KIF17 do not localize to neuronal primary cilia. Scale bars represent 100 µm for A and 10 µm B-E.

**Figure S2.**
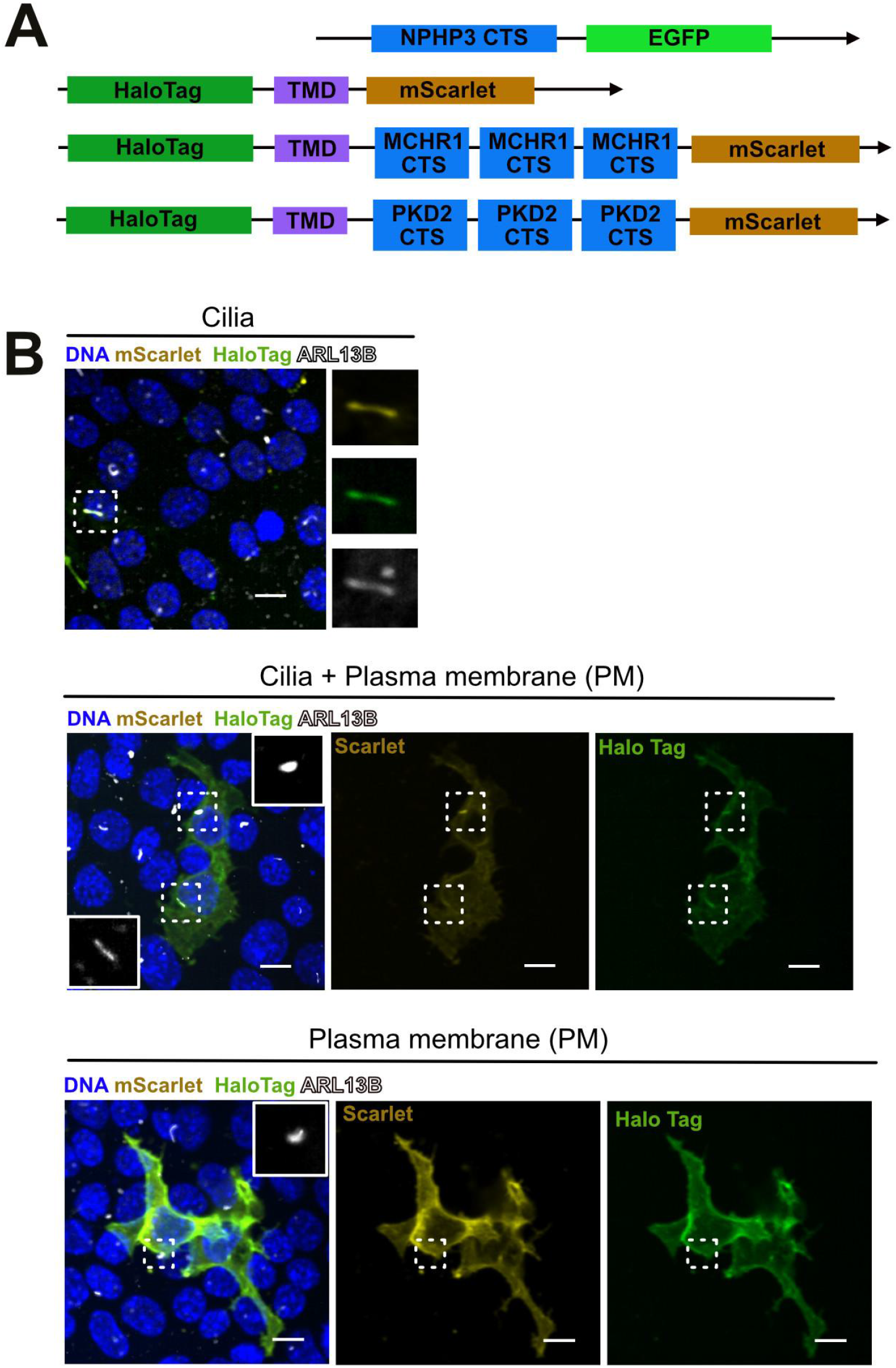
Cilia targeting construct design and their localisation in hiPSC-derived cells. **A)** Schematic design of the Cilia-APEX construct (top), TM control construct lacking ciliary localization sequences, and constructs with CTSs from MCHR1 and PKD2. **B)** Representative images of cells with exclusively ciliary localisation, mixed ciliary and non-ciliary localisation, or exclusively non-ciliary localisation. Scale bars represent 10 µm.

**Figure S3.**
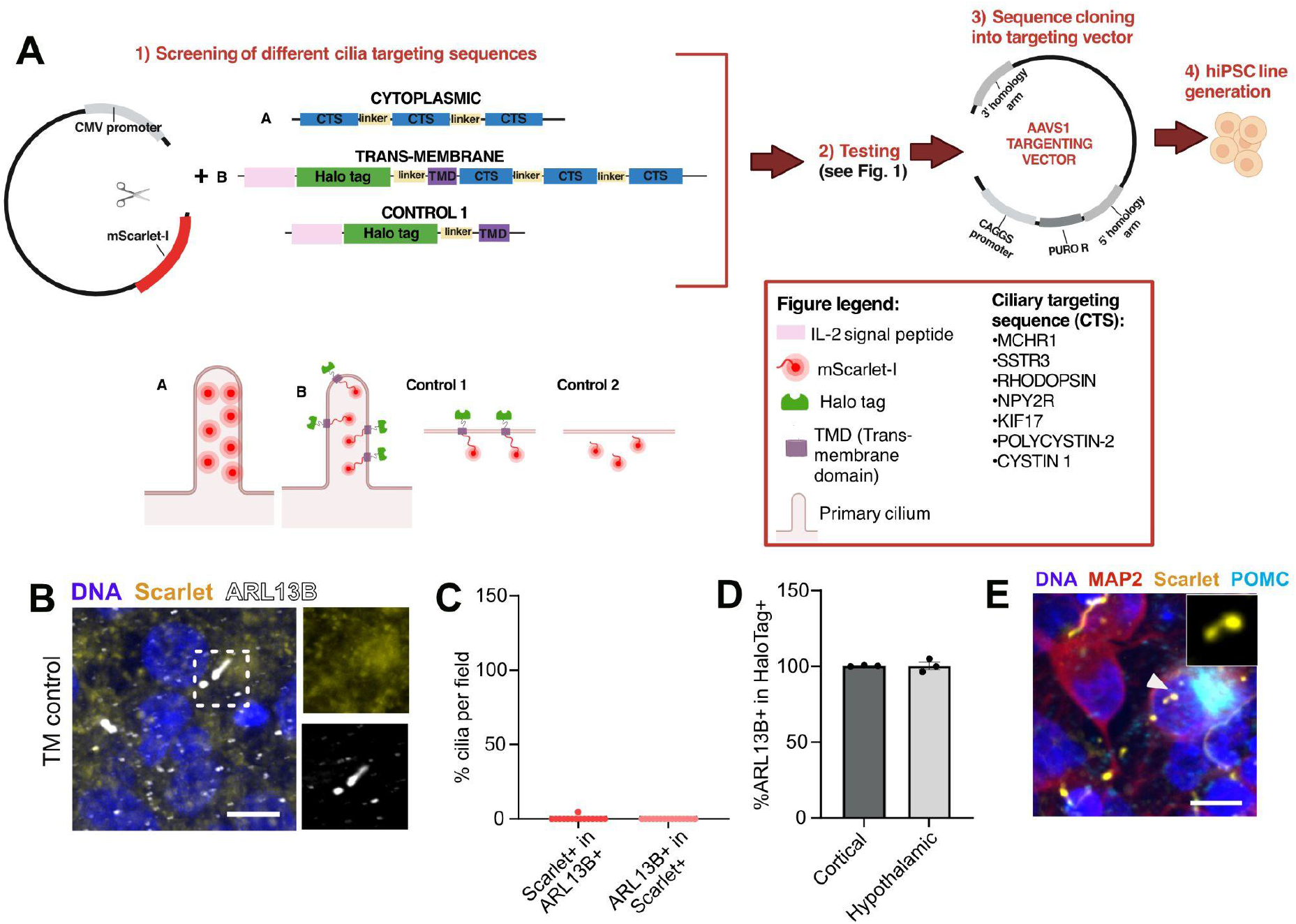
Design and testing of ciliary reporter cell lines. **A)** Experimental schematic for 1) screening different cilia targeting sequences by transient transfection of expression plasmids driven by a CMV promoter and visualized by mScarlet, followed by 2) testing by transient transfection, 3) cloning sequences into an AAVS1 targeting vector with positive and negative selection, and 4) generation of stable hiPSC lines. **B)** The stable hiPSC line carrying the TM control plasmid that lacks CTSs does not show mScarlet co-localisation with ARL13B. **C)** Quantification of ARL13B+ structures expressing mScarlet in cells transfected with the TM control plasmid, n = 311 ARL13B+ structures. **D)** Quantification of HaloTag-expressing structures that co-express ARL13B in hiPSC-derived cortical and hypothalamic cells. N = 3 technical replicates, n = 25 +/- 3 (hypothalamic) and 266 +/- 55 (cortical) HaloTag+ structures per replicate. **E)** Human iPSC-derived hypothalamic neurons immunopositive for pro-opiomelanocortin (POMC) have primary cilia that are labeled by endogenous mScarlet fluorescence. Scale bars represent 10 µm in B and E.

**Figure S4.**
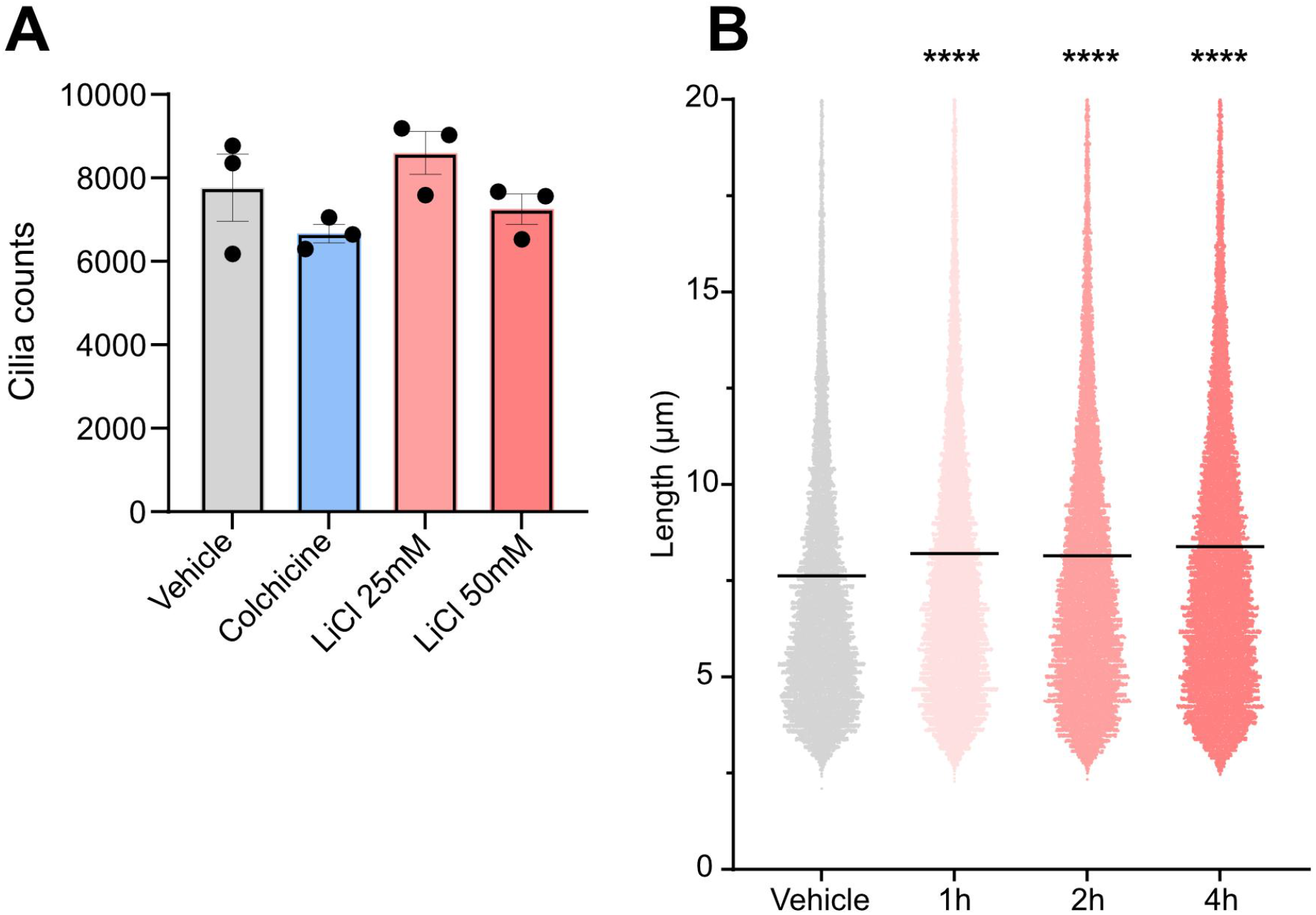
Effects of LiCl on primary cilia length in human neurons. **A)** Total cilia numbers quantified across parallel cultures of hiPSC-derived hypothalamic neurons and treated with colchicine or LiCl are similar. B) Neuronal cultures treated with a vehicle control or 25 mM LiCl for 1, 2, 4, or 24 hours show time-dependent changes in primary cilia length relative to controls. N = 3 technical replicates per condition, and 8328 +/- 720 quantified structures per replicate. ****p<0.0001 by Kruskal-Wallis test.

